# cancerAlign: Stratifying tumors by unsupervised alignment across cancer types

**DOI:** 10.1101/2020.11.17.387860

**Authors:** Bowen Gao, Yunan Luo, Jianzhu Ma, Sheng Wang

## Abstract

Tumor stratification, which aims at clustering tumors into biologically meaningful subtypes, is the key step towards personalized treatment. Large-scale profiled cancer genomics data enables us to develop computational methods for tumor stratification. However, most of the existing approaches only considered tumors from an individual cancer type during clustering, leading to the overlook of common patterns across cancer types and the vulnerability to the noise within that cancer type. To address these challenges, we proposed cancerAlign to map tumors of the target cancer type into latent spaces of other source cancer types. These tumors were then clustered in each latent space rather than the original space in order to exploit shared patterns across cancer types. Due to the lack of aligned tumor samples across cancer types, cancerAlign used adversarial learning to learn the mapping at the population level. It then used consensus clustering to integrate cluster labels from different source cancer types. We evaluated cancerAlign on 7,134 tumors spanning 24 cancer types from TCGA and observed substantial improvement on tumor stratification and cancer gene prioritization. We further revealed the transferability across cancer types, which reflected the similarity among them based on the somatic mutation profile. cancerAlign is an unsupervised approach that provides deeper insights into the heterogeneous and rapidly accumulating somatic mutation profile and can be also applied to other genome-scale molecular information.

**Availability:** https://github.com/bowen-gao/cancerAlign

## 1 Introduction

Tumor stratification aims at dividing a heterogeneous collection of tumors into biologically meaningful and clinically actionable subtypes based on the similarity of molecular profiles, such as gene expression and somatic mutation[1]. Owing to the substantial heterogeneity of cancers, tumor stratification paves the path for personalized treatment, moving away from the conventional one-size-fits-all approach[2]. To better understand cancer heterogeneity, large-scale cancer genomics projects, such as the Cancer Genome Atlas (TCGA), the International Cancer Genome project, and the Memorial Sloan Kettering-Integrated Mutation Profiling project have systematically profiled thousands of tumors, holding the promise to realize personalized treatment[3–6]. Among the collected genome-scale omics data, somatic mutation profiles have been used to discover causal drivers of tumors[7–13] and further reveal informative cancer subtypes[14,15]. In pursuit of this vision, computational approaches have been developed to stratify tumors according to high-dimensional, noisy and sparse somatic mutation profiles[1,16–21].

Existing computational approaches mainly exploited two machine learning techniques to advance tumor stratification: dimensionality reduction[22–27] and network-based aggregation of individual mutations[1,17–19,28,29]. For example, NBS used molecular networks to aggregate individual gene mutation into higher level functions and structures in cancer cells[1]. Wang et al. further extended this framework by exploiting mutual exclusivity to construct a stringent molecular network for propagation[16]. Despite the encouraging performance of these methods, none of them has considered analyzing tumor samples from multiple cancer types simultaneously, inevitably overlooking shared mutation patterns across cancer types. Such mutation patterns, including network modules and signaling pathways, have been discovered through pan-cancer analysis, and further reveal the similarities and differences across cancer types[30]. Moreover, signals from mutations that are rare in one cancer type might be boosted from functionally similar mutations from other cancer types.

Nevertheless, jointly analyzing tumors from two or more cancer types is challenging due to the incomparable somatic mutation profiles across cancer types. Tumors are wildly heterogeneous across cancer types, and different cancer types are known to be driven by different genes and pathways[31]. Simply selecting features of common genes across cancer types might produce clusters that are biased to across-cancer-type signals rather than within-cancer-type signals. A plausible solution is to learn a nonlinear mapping between mutation profiles of two cancer types[32]. The nonlinear mapping does not presume that two cancer types share the exact same driver genes or pathways, thus better modeling the heterogeneous across cancer types. However, supervised alignment approaches cannot be adopted to learn this mapping due to the lack of aligned tumor samples across cancer types.

In this paper, we propose cancerAlign, an unsupervised approach to stratify tumors through learning tumor alignment across cancer types. The key idea of cancerAlign is to learn a shared latent space between the somatic mutation profile of a target cancer type and a source cancer type. Tumors of the target cancer type are then clustered in this shared latent space so that common patterns across cancer types can be exploited to assist the stratification. Due to the absence of annotated aligned tumor samples across cancer types, cancerAlign learns the mapping at the population level using adversarial learning. Moreover, cancerAlign automatically selects source cancer types according to the silhouette score between tumor samples in the shared latent space and tumor samples in the original space. We evaluated cancerAlign on 7,134 tumors spanning 24 cancer types from TCGA[33]. cancerAlign obtained substantial improvement against the method that only considers a single cancer type in both tumor stratification and cancer gene prioritization. Our method further reveals the similarity and transferability across cancer types, presenting new opportunities for personalized treatment and drug repurposing[34].

## 2 Problem definition

### 2.1 Problem definition of conventional tumor stratification

The input of tumor stratification is a binarized somatic mutation matrix 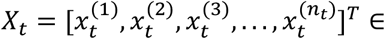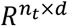 of a particular cancer type *c_t_*, where *n_t_* is the number of tumors in the dataset, and *d* is the number of genes. The goal is to cluster these *n_t_* tumors into *k* clusters based on the binarized somatic mutation matrix. The output is the cluster label vector *l_t_* ∈ *R*^*n*^*t*, which would be evaluated by the difference of survival rates among clusters.

### 2.2 Problem definition of cancerAlign

Different from conventional tumor stratification methods that only consider a single cancer type, cancerAlign leverages multiple source cancer types to assist the clustering of the target cancer type. In particular, cancerAlign consists of a mapping step and a clustering step. In the mapping step, the input is the binarized somatic mutation matrices of *m* source cancer types 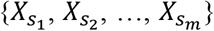 where 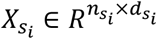 and the target cancer types *X_t_* ∈ *R*^*n*_*t*_×*d*_*t*_^. We aim at learning a mapping 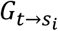 from the target cancer type *t* to each source cancer type *s_i_*. For patient *j* in the target cancer type *t*, 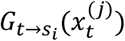 is in the same feature space as 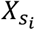. The output of the mapping step is then a mapped matrix 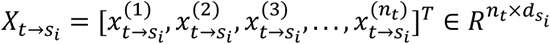. In the clustering step, cancerAlign clusters tumors using the mapped mutation matrix *X*_*t*→*s*_ instead of the original matrix *X*_*t*_. cancerAlign then obtains a cluster label vector 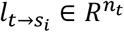 by mapping to each source cancer type *s_i_*. We then integrate these cluster label vectors to obtain the final cluster label vector *l_t_* ∈ *R*^*n*_*t*_^.

## 3 Methods

Supervised approaches cannot be used to align tumors between two cancer types due to the lack of annotated aligned tumors. The incomparable feature space among different cancer types also poses difficulty to jointly analyze them. To address these problems, cancerAlign uses an adversarial learning method to perform unsupervised alignment between two cancer types, which does not require aligned samples or same features space between two cancer types. Tumors are then clustered in this aligned space and cluster label vectors from multiple source cancer types are further integrated to provide a robust clustering result (**Fig. 1**).

**Fig. 1.**
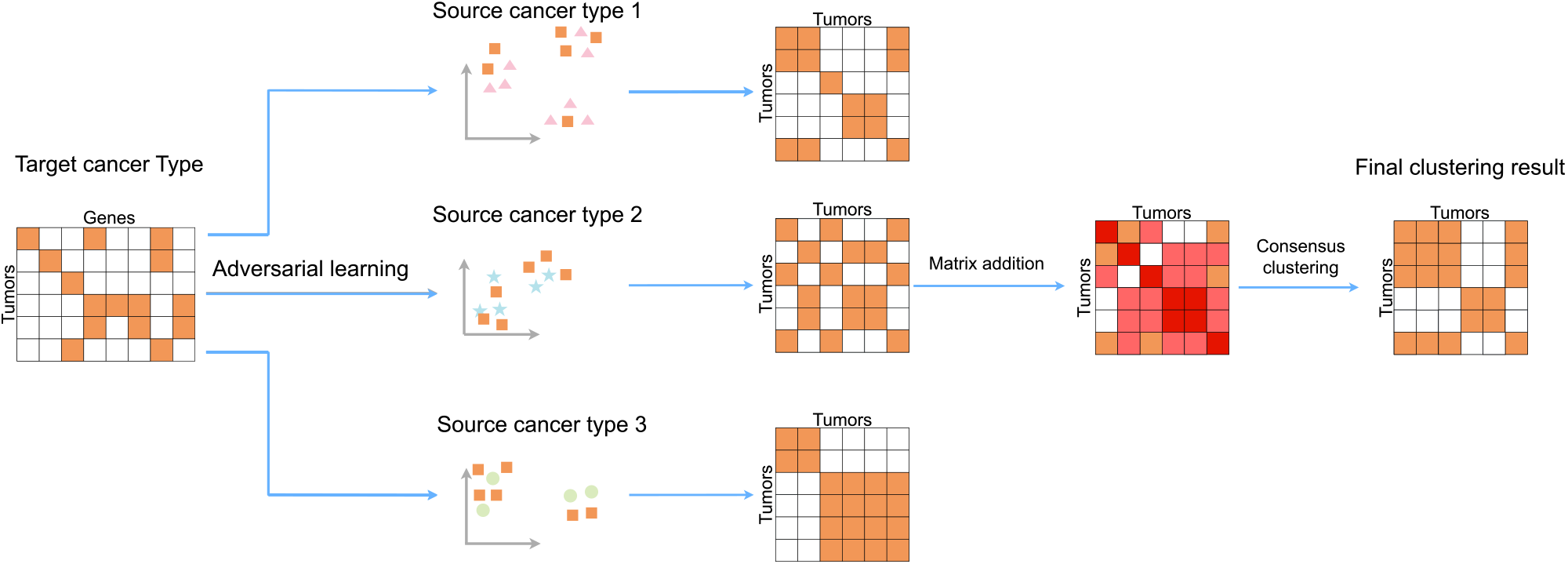
Flowchart of cancerAlign. cancerAlign first maps the somatic mutation profile of tumors from the target cancer type to different source cancer types. It then clusters these tumors in the latent space of each source cancer type. As a result, cancerAlign obtains an individual cluster label vector for each source cancer type. These cluster label vectors are then aggregated together by consensus clustering to get the final tumor stratification result.

### 3.1 Unsupervised alignment via adversarial learning

We proposed to use adversarial learning to map tumor samples between cancer types. Adversarial learning has been widely used in various machine learning fields, such as computer vision[35], natural language processing[36], and computational biology[37]. Our adversarial learning framework aims at training a generator and a discriminator. The generator aims at generating corresponding samples in the source caner type given a sample in the target cancer type. The discriminator aims at classifying samples in the source cancer type space into generated samples or real samples. A good generator should make the classification of the discriminator difficult, while a good discriminator would make the generation of samples more challenging. Consequently, the generator and the discriminator iteratively advance each other in a competition mode, and the resulsted generator would be used to map tumors from the target cancer type to the source cancer type.

The generator is the mapping function 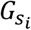 consisting of an encoder and a decoder. Given the input data 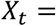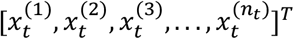 of a particular cancer type *c_t_*, where each 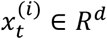 representing the feature vector of tumor *i*. The encoder transforms the original input vector 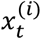 into a low-dimensional vector 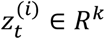. Then this low-dimensional vector is used by the decoder to generate the output vector 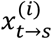 that shares the same feature space as cancer type *s*. The output matrix by mapping tumors in cancer type *t* to cancer type *s* is thus 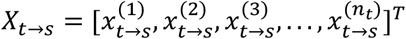.

We also trained a discriminator *D* to discriminate between feature vectors from *X*_*S*_ and from *X*_*t*→*S*_. Given the input data 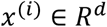, the discriminator is a binarized classifier *D*: *R*^*d*^ → {0,1}, where 1 means that *x*^(*i*)^ is from *X*_*S*_ and 0 means that *x*^(*i*)^ is from *X*_*t*→*S*_.

The discriminator *D* is optimized to correctly classify whether the input tumor sample is from *X*_*S*_ or *X*_*t*→*S*_, while the generator *G* is optimized to generate tumor samples that can fool the discriminator. Thus the discriminator and the generator form a two-player minimax game with a value function *V*(*D*, *G*):

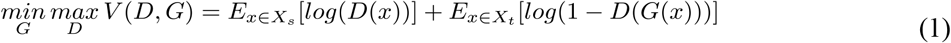

In the training stage, we updated the parameters of the generator and the discriminator iteratively. Each one has its own loss function.

#### Discriminator objective

Let θ_*D*_ be the parameters of the discriminator. The learning objective of the discriminator is to successfully decide whether the input vector is from the source cancer type, or generated by the mapping function. We treated 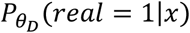 as the probability that a vector *x* is from the real source cancer, rather than the generator. Then the loss for discriminator is:

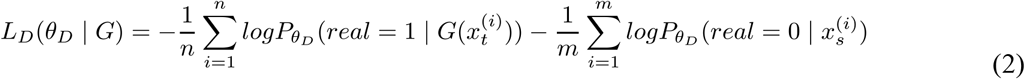

#### Generator objective

The learning objective for the generator is to fool the discriminator. We wanted to train the *G* such that the discriminator cannot correctly classify whether the input tumor sample is from *X*_*S*_ or *X*_*t*→*S*_. Then the loss for the generator is:

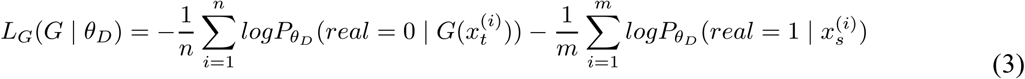

### 3.2 Selecting source cancer types

Our method considered multiple source cancer types and learnt a mapping function from target cancer type to each source cancer type. However, due to the heterogeneity across cancer types, many source cancer types may not align well with the given target cancer type, thus introducing additional noise to the clustering step. To increase the signal-to-noise ratio, we proposed to only consider source cancer types that are indistinguishable with the target cancer type after the mapping.

In particular, we used the Silhouette score to measure whether tumors in the mapped target cancer space are indistinguishable with tumors from the original cancer type space. Silhouette score has been widely used to evaluate the accuracy of clustering results[38,39]. A larger score indicates a better clustering result. We first treated *X*_*t*_ and *X*_*t*→*S*_ as two different clusters, then calculated the Silhouette score between these two clusters, corresponding to the similarity between tumor samples in the mapped target cancer space and tumor samples in the original cancer type space. Source cancer types that had closer to zero Silhouette scores were in favor since it indicated that the generated feature vectors of cancer type *X*_*t*→*S*_ was indistinguishable from the target cancer type *X*_*t*_. Since there were multiple source cancer types, we only considered the top five cancer types that had the closest to zero Silhouette scores for the next step consensus clustering.

Due to the lack of a single objective loss function, it is challenging to know how well a generator has been trained[40]. The discriminator and the generator are trained in a competitive mode, which further makes it difficult to determine convergence. We selected the number of iterations based on the number of patients in the smallest cluster. Let *q_e_* be the number of patients in the smallest cluster based on the cluster label vector in the epoch *e*. We found that *q_e_* tends to become substantially small (e.g., less than 5) after a large number of iterations by iteratively training the generator and the discriminator. A well-known issue in the adversarial training is model collapse, which means the generator producing similar samples (partial collapse) or in the worst case repetitively generating the same sample (complete collapse)[41]. In this case, we regarded such a small *q_e_* as the signal of the model collapse of GAN[42] and leading to trivial clusters. In order to avoid the trivial cluster results, we used the epoch that has the largest *q_e_*.

### 3.3 Consensus clustering across selected source cancer types

We used K-means as the base clustering algorithm to cluster tumors based on the mapped mutation profile *X*_*t*→*S*_. Due to the high dimensionality of the feature vectors, we first used principal component analysis (PCA) to reduce the dimensionality before clustering. By considering different source cancer types, we could obtain multiple cluster label vectors. We then aggregated these different label vectors using consensus clustering to obtain the final cluster label vectors. Consensus clustering assesses the consensus among multiple existing clustering results to obtain a more robust clustering result[43]. Specifically, we first obtained individual cluster label vector 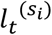 for clustering tumors of cancer type *t* in the feature space of each source cancer type 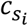. Then we used the label vector to get the connectivity matrix *CM*^(*S*_*i*_)^. Each *CM*^(*S*_*i*_)^ is an *n_t_* × *n_t_* matrix, with rows and columns representing patients of the target cancer type. The entries in this matrix are defined as:

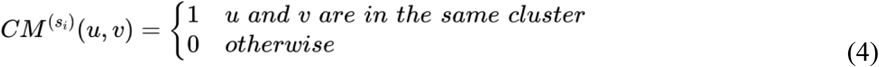

Then we defined the consensus matrix *W*, where each entry (*u*, *v*) inside the consensus matrix represents the frequency that patient *u* and patient *v* are within the same clusters according to 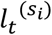.

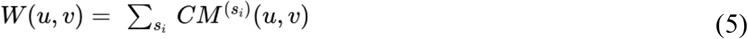

We used *W* as our new feature matrix for K-means to obtain the final clustering result.

## 4 Results

### 4.1 Somatic mutation and patient survival data

We downloaded somatic mutation profiles of tumors in The Cancer Genome Atlas (TCGA) from GDAC Firehose website (http://gdac.broadinstitute.org, 11th February 2016). In total, we collected somatic mutation profiles of 7,134 tumors belonging to 24 different cancer cohorts, including BRCA, BLCA, CESC, CHOL, COAD, DLBC, GBM, HNSC, KICH, KIRC, LGG, LIHC, LUAD, LUSC. OV, PAAD, PRAD, READ, SARC, STES, TFCT, THCA, UCEC, UVM. These cancer types contain different numbers of tumors, ranging from 35 to 974, with an average of 297 tumors. The number of genes in each cancer type is between 1,305 and 16,859, with an average of 10,234 genes. There are in total 18,022 unique genes across all 24 cancer types. We also obtained the survival data from the TCGA dataset for these tumors. We used the survival data and known cancer genes to evaluate the performance of the tumor stratification. The known cancer genes we used are obtained from Iorio et al.[44]. It contains cancer genes for different cancer types, and only high confidence genes are considered. Those cancer types have different numbers of cancer genes, ranging from 8 to 62, with an average of 29.33 cancer genes.

### 4.2 Experimental setting

To train our adversarial learning framework, we followed the standard training process from previous work[45]. We made samples in each mini-batch either all from the target cancer type, or all from the mapped target cancer type. As a result, no batch contained mixed samples. We trained the model for 100 epochs. For the Adam optimizer, we set the starting learning rate, the exponential decay rate for the first moment estimates, and the exponential decay rate for the second moment estimates to 2e-5, 0.9, 0.999, respectively.

We compared cancerAlign with two comparison approaches. The first one, named as **clustering without alignment**, is the conventional tumor stratification on an individual cancer type. K-means clustering algorithm is directly applied to the original binarized data of target cancer type (i.e., *X*_*t*_). We compared our method with clustering without alignment to study the importance of jointly considering multiple cancer types. The second one, named as **Pan-Cancer clustering**, combined tumors from all cancer types and then performed the clustering. In particular, given the target cancer type *c_t_* that we aimed to stratify and another source cancer type 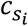, we jointly clustered tumors from these two cancer types using common genes between *c_t_* and 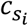 as features. We then extracted the cluster label vector of tumors in *c_t_* and applied the consensus clustering method described in 3.3 to ensemble different cluster label vectors generated by using different 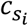. We compared our method with Pan-Cancer clustering to investigate the effect of learning a nonlinear mapping between cancer types.

We evaluated the clustering performance based on survival rates among different clusters. We used the log-rank test[46], where a small *p*-value indicates that the survival rates are significantly different among clusters. Let the number of clusters be *k*. We thoroughly examined different *k* across from 2 to 6. For each target cancer type, we also selected the best results from experiments of different *k*. We also investigated whether our model can identify cancer genes by comparing differentially mutated genes to a list of known cancer genes. We used the chi-squared test to identify cancer genes based on the cluster label vectors from our method. Specifically, for each gene *i* in cancer type *t*, we obtained two equal-sized vectors. The first vector is the output of cancerAlign 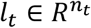, where each entry represents the cluster label of a tumor. The second vector is the mutation vector *g*_*i*_, where 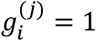 if gene *i* is mutated in tumor *j*, otherwise 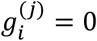. We then applied chi-squared test on *l*_*t*_ and *g*_*i*_ and obtained a *p*-value for gene *i*. We created the cancer genes list of our method according to these *p*-values. AUROC was used to evaluate the performance of cancer gene identification.

To investigate which source cancer type *s* is most helpful to the target cancer type *t* in tumor stratification, we directly clustered tumors using each *X*_*t*→*S*_ without performing consensus clustering. To find novel cancer genes, we first selected cancer types that have the good AUROC scores using our method. For each cancer type, we identified the top 10 genes with the lowest *p*-values and determined the ones that are not in the known cancer gene list as novel cancer genes. To map across more than two cancer types, we sequentially considered these cancer types. For example, to map cancer type *t* to *s* and then to *r*, we first learnt a mapping between *X*_*t*_ and *X*_*S*_ and obtained the mapping *G*_*t*→*S*_. We then learnt another mapping between *X*_*S*_ and *X*_*r*_. Tumors in cancer type *t* were finally clustered using *G*_*t*→*S*→*r*_(*X*_*t*_).

### 4.3 cancerAlign improves tumor stratification

We first sought to investigate whether cancerAlign can improve tumor stratification by using source cancer types to assist the clustering of tumors in the target cancer type. To this end, we compared cancerAlign with clustering without alignment on 24 cancer types across the number of cluster *k* from 2 to 6. We found that cancerAlign substantially outperformed clustering without alignment on a large number of cancer types (**Fig. 2a,b,c**). Overall, cancerAlign obtained significant clustering for 8 cancer types, whereas clustering without alignment only has significant clustering for 6 cancer types (*p*-value<0.01 log-rank test) (**Fig. 2a**). Moreover, cancerAlign obtained improved clustering results for 8 out of 10 cancer types which obtained significant *p*-value by at least one method. For example, cancerAlign obtained a log-rank survival association *p*-value less than 4e-4 for KIRC when *k* = 3, which is much more significant than the *p*-value of 0.2 by clustering without alignment (**Fig. 2b, Fig. 3a**). The consistent superior performance of cancerAlign across different numbers of clusters demonstrates the effectiveness of using source cancer types to assist the clustering of a specific target cancer type.

**Fig 2.**
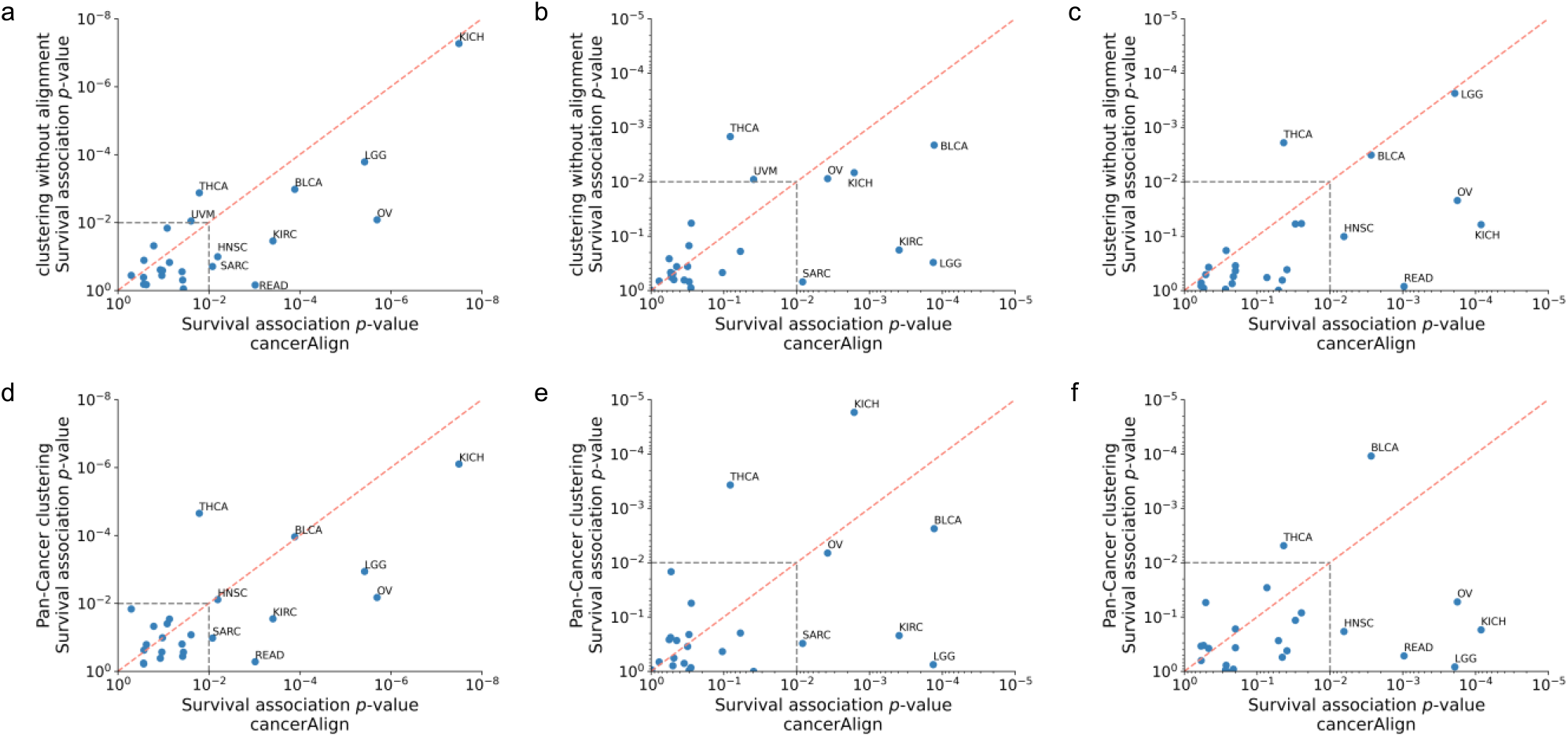
Comparison between our method and comparison approaches on tumor stratification. Each dot represents one of 24 cancer types. x-axis shows the survival association *p*-value by using our method, while y-axis shows the survival association *p*-value of the comparison approach. **a,b,c,** Scatter plot showing the comparison of our method with clustering without alignment using the best clustering result across *k* from 2 to 6(a), *k*=3(b), *k*=5(c). **d,e,f,** Scatter plots showing the comparison of our method with Pan-Cancer clustering using the best clustering result across *k* from 2 to 6(d), *k*=3(e), *k*=5(f).

**Fig 3.**
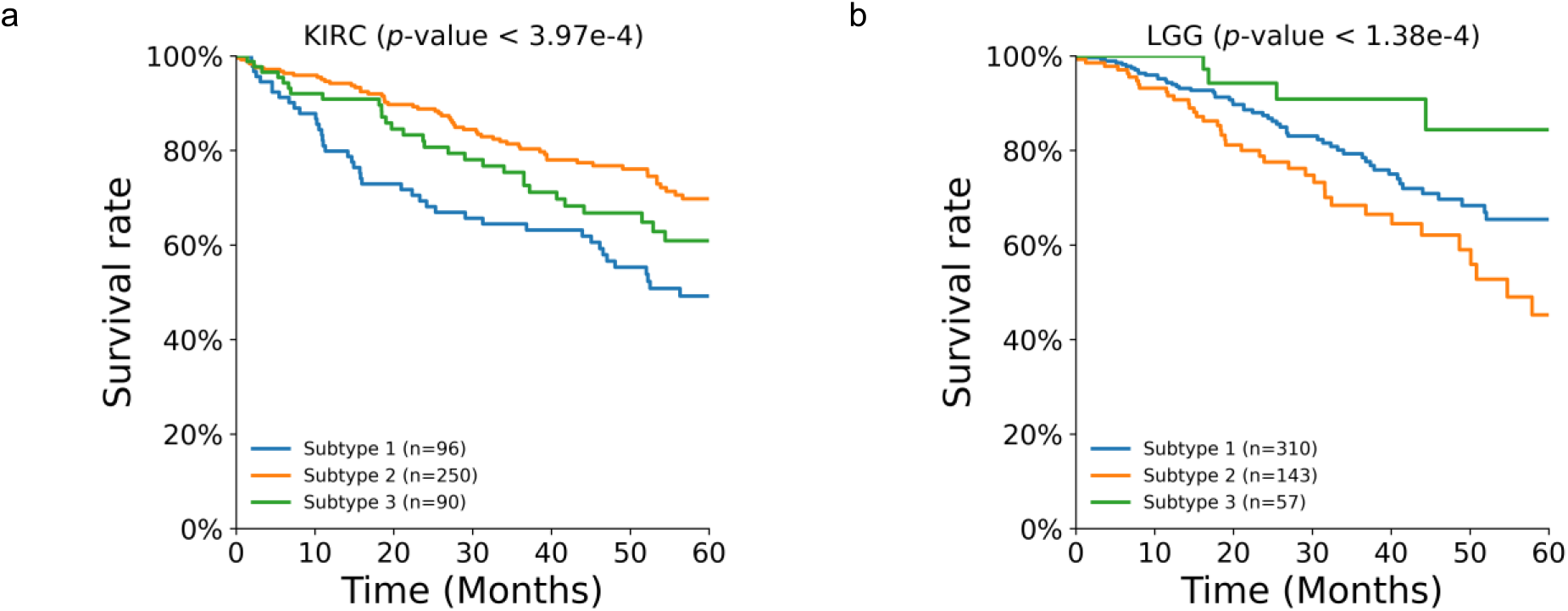
Kaplan-Meier (KM) survival plots using cancerAlign. **a,** KM survival plots for KIRC using cancerAlign when the number of clusters equal to 3. **b,** KM survival plots for LGG with cancerAlign when the number of clusters equal to 3.

Next, we compared cancerAlign with Pan-Cancer clustering, which also clustered tumors using multiple cancer types. We found that cancerAlign also greatly improved Pan-Cancer clustering (**Fig. 2d,e,f**). Among 24 cancer types, cancerAlign obtained a significant *p*-value for 8 cancer types, which is still larger than the 6 cancer types of Pan-Cancer clustering (**Fig. 2d**). Out of the 9 cancer types that obtained significant *p*-value by at least one of the methods, cancerAlign achieved improved clustering results on 7 of them. For instance, cancerAlign obtained a log-rank survival association *p*-value<2e-4 for LGG when *k* = 3, which is substantially better than the *p*-value of 0.9 by Pan-Cancer clustering (**Fig. 2f, Fig. 3b**). The superior performance of cancerAlign indicates the importance of learning a nonlinear mapping to align two cancer types rather than a simple combination between them. We further noticed that the performance of Pan-Cancer clustering is in general better than clustering without alignment, which raises our confidence about leveraging other cancer types to assist tumor stratification.

### 4.4 cancerAlign improves cancer gene identification

Next, we studied whether our method can also improve the cancer gene identification. We first compared cancerAlign with clustering without alignment on cancer gene identification for 24 cancer types (**Fig. 4a**). We observed that cancerAlign achieved AUROCs greater than 0.80 for 8 cancer types, whereas none of the 24 cancer types had an AUROC larger than 0.80 by using clustering without alignment. Our method achieved an average of 0.74 AUROC, which is 40% better than the 0.53 AUROC of clustering without alignment. We then compared cancerAlign with Pan-Cancer clustering and again observed that cancerAlign outperformed the Pan-Cancer clustering method in cancer gene identification (**Fig. 4b**). All AUROCs of 24 cancer types were less than 0.80 by using Pan-Cancer clustering. In contrast, 8 out 24 cancer types had an AUROC greater than 0.80 by using cancerAlign. Overall, cancerAlign Pan-Clustering obtained an average of 0.59 among all cancer types, which is much lower than the 0.74 AUROC value achieved by our method. The superior performance against two comparison approaches demonstrates the effectiveness of using other cancer types to assist the cancer gene identification of a specific cancer type.

**Fig 4.**
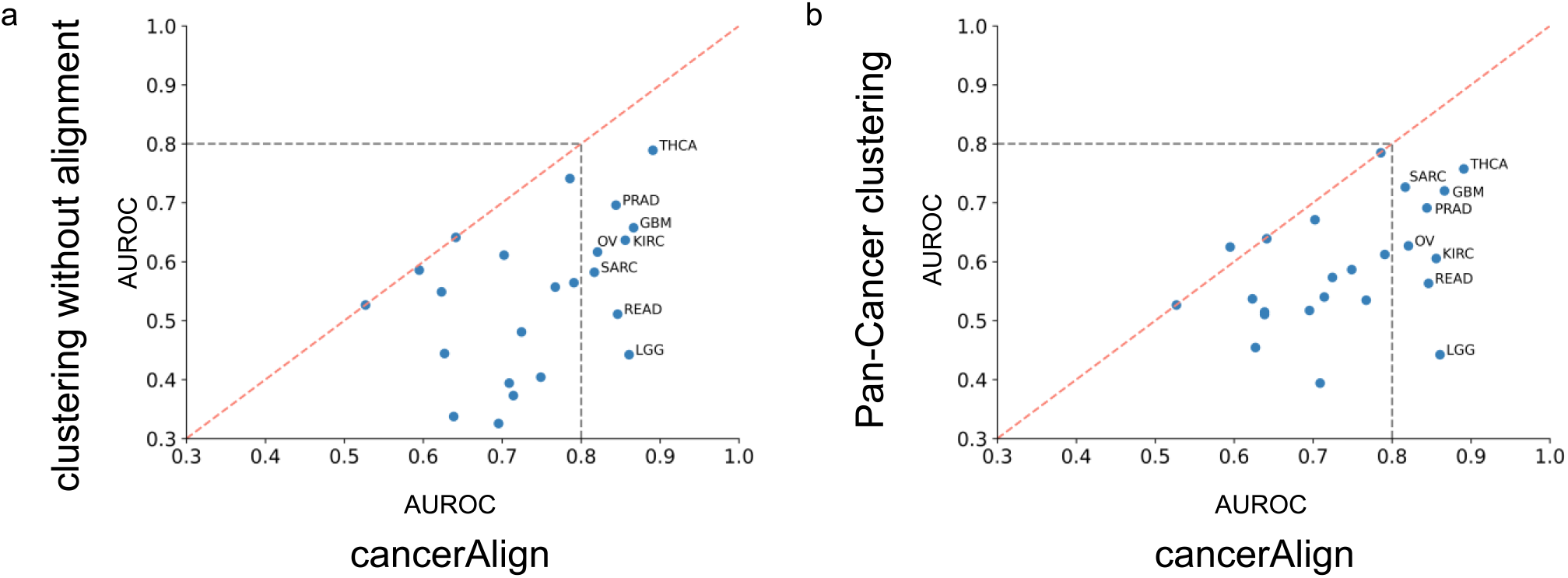
Comparison between our method and comparison approaches on cancer gene identification. Each dot represents one of 24 cancer types. x-axis shows the AUROC by using our method, while y-axis shows the AUROC of the comparison approach. **a,** Scatter plot showing the comparison between our method and clustering without alignment. **b,** Scatter plot showing the comparison between our method and Pan-Cancer clustering.

We then examined the novel cancer genes identified by cancerAlign and found that many of them can be verified by existing literature. For example, cancerAlign found that TTN is important for tumor straticiotan in READ(*p*-value<2e-06), which is not in the known cancer gene list. A recent study showed that the mutation count of TTN had a large correlation coefficient with the tumor mutation burden, which is an emerging biomarker in colorectal cancer[47]. TTN mutation was enriched in samples possessing high immunostimulatory signatures, partially reflecting the association between large mutation load and high TMB status. We also found that CALN1 might be a novel cancer gene for GBM, which obtained a *p*-value<0.01 by our method. Lu et al. found that GBM patients with overexpressed CALN1 had better overall survival, which suggests that CALN1 might be a potential biomarker for identifying high-risk patients with GBM[48].

### 4.5 cancerAlign identify transferable cancer types

Motivated by the improved performance of cancerAlign in tumor stratification and cancer gene identification, we then examined which two cancer types could produce a good mapping that leads to superior tumor stratification. We summarized the cancer type pairs that have the best tumor stratification result in **Table 1**. We observed a great improvement of our method on these cancer types. For example, by mapping Kidney Renal Clear Cell Carcinoma (KIRC) to Kidney Chromophobe (KICH), cancerAlign obtained a *p*-value of 3e-4, which is more significant than the comparison approach. We found that some of these cancer type pairs can be supported by literature. For example, a recent immune genes analysis[49] showed that LGG and KIRC had the largest numbers of prognostic immune genes (PIGs) among 22 analyzed cancer types. Moreover, they found that the number of risk PIGs (hazard ratio > 1) was significantly higher than that of protective PIGs for both KIRC and LGG. By learning a nonlinear mapping between cancer types, cancerAlign identified transferable cancer types, which presents new opportunities for cancer drug development and repurposing.

**Table 1.** Cancer type pairs that have the most prominent tumor stratification improvement using cancerAlign. Given a target cancer type, the source cancer type that leads to the best tumor stratification result is shown. The *p*-values of our method and clustering without alignment are also shown.

| Target cancer | Best source cancer | Number of clusters | $P$-value of comparison approach | $P$-value of our method |
| --- | --- | --- | --- | --- |
| UVM | LUSC | 4 | 3.60E-01 | 1.11E-06 |
| LGG | LUSC | 4 | 1.62E-01 | 1.11E-06 |
| LGG | KIRC | 3 | 3.05E-01 | 2.76E-06 |
| BLCA | OV | 3 | 2.11E-03 | 8.59E-05 |
| KIRC | KICH | 3 | 1.80E-01 | 3.09E-04 |
| OV | GBM | 3 | 8.77E-03 | 3.49E-04 |
| HNSC | GBM | 6 | 1.81E-01 | 5.32E-03 |
| SARC | KICH | 3 | 6.95E-01 | 6.06E-03 |

Finally, we explored whether our method can be used to sequentially align more than two cancer types. As a proof-of-concept, we illustrated two examples where sequentially aligning three cancer types had better performance than only aligning two cancer types (**Table 2**). For example, mapping SARC to BRCA can improve the tumor stratification result from *p*-value of 1.96e-1 to *p*-value of 1.88e-2 when clustering tumors into 4 subtypes. By further mapping from BRCA to COAD, the tumor stratification result is further enhanced to a *p*-value of 4.62e-3. In HNSC, cancerAlign improved the clustering result by mapping HNSC to LGG and obtained a *p*-value of 5.17e-3. By further mapping these tumors to LUSC, the *p*-value improved to 6.14e-4. The observed improvement of sequentially mapping multiple cancer types suggests the possibility to further improve cancerAlign by automatically determining the cancer mapping sequence and provide novel insights into cancer research.

**Table 2.**
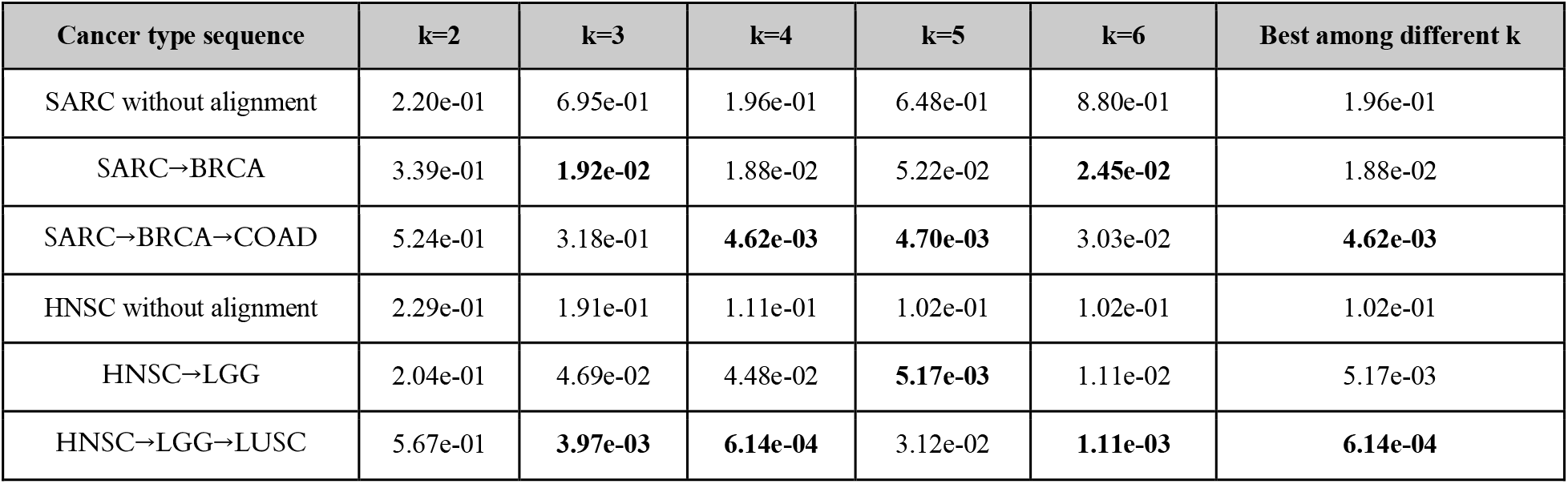
Comparison among clustering without alignment, clustering by aligning two cancer types, and clustering by aligning three cancer types on tumor stratification. We show different results for k (number of clusters) from 2 to 6, and the best clustering results for k across 2 and 6. Best and significant *p*-values for each number of cluster are shown in bold.

| Cancer type sequence | k=2 | k=3 | k=4 | k=5 | k=6 | Best among different k |
| --- | --- | --- | --- | --- | --- | --- |
| SARC without alignment | 2.20e-01 | 6.95e-01 | 1.96e-01 | 6.48e-01 | 8.80e-01 | 1.96e-01 |
| SARC→BRCA | 3.39e-01 | **1.92e-02** | 1.88e-02 | 5.22e-02 | **2.45e-02** | 1.88e-02 |
| SARC→BRCA→COAD | 5.24e-01 | 3.18e-01 | **4.62e-03** | **4.70e-03** | 3.03e-02 | **4.62e-03** |
| HNSC without alignment | 2.29e-01 | 1.91e-01 | 1.11e-01 | 1.02e-01 | 1.02e-01 | 1.02e-01 |
| HNSC→LGG | 2.04e-01 | 4.69e-02 | 4.48e-02 | **5.17e-03** | 1.11e-02 | 5.17e-03 |
| HNSC→LGG→LUSC | 5.67e-01 | **3.97e-03** | **6.14e-04** | 3.12e-02 | **1.11e-03** | **6.14e-04** |

## 5 Conclusion and discussion

In this paper, we have presented cancerAlign, a novel computational method that stratifies tumors through jointly aligning multiple cancer types. We used adversarial learning to map tumors from the target cancer type to multiple source cancer types. We then applied consensus clustering to integrate clustering based on different source cancer types. We observed substantial improvement against clustering without alignment and Pan-Cancer clustering in tumor stratification and cancer gene prioritization.

Our method is inspired by the recent progress in unsupervised machine translation and single cell integration. Conventional machine translation relied on supervised training using parallel data between two languages. Recently, unsupervised machine translation has become feasible by exploring the shared latent space across word-occurrence patterns across languages[36]. In addition to also performing unsupervised alignment between cancer types, we further integrated information from multiple source cancer types and proposed to select source cancer types based on clustering agreement. Another line of the exciting progress is in single cell integration, where methods such as Scanorama[50] have been proposed to integrate single cell datasets without known aligned samples. We found that adversarial learning resulted in better alignment, partially due to the incomparable feature spaces among cancer types.

While cancerAlign obtained substantial improvement here, there are still several potential future directions we would like to investigate. Currently, we only considered somatic mutation data. We plan to jointly align somatic mutations with other genomics data such as gene expression profiles. Other than tumors, cell lines[44] and patient-derived tumor xenograft[51] models are also in pressing needs for cancer research. We would also like to incorporate them into our current framework. Finally, molecular networks might further advance cancerAlign through grouping sparse mutations into high-level modules[1].

## Notes

### Competing Interest Statement

The authors have declared no competing interest.

